# Functional assessment of cell entry and receptor usage for lineage B β-coronaviruses, including 2019-nCoV

**DOI:** 10.1101/2020.01.22.915660

**Authors:** Michael Letko, Vincent Munster

## Abstract

Over the past 20 years, several coronaviruses have crossed the species barrier into humans, causing outbreaks of severe, and often fatal, respiratory illness. Since SARS-CoV was first identified in animal markets, global viromics projects have discovered thousands of coronavirus sequences in diverse animals and geographic regions. Unfortunately, there are few tools available to functionally test these novel viruses for their ability to infect humans, which has severely hampered efforts to predict the next zoonotic viral outbreak. Here we developed an approach to rapidly screen lineage B betacoronaviruses, such as SARS-CoV and the recent 2019-nCoV, for receptor usage and their ability to infect cell types from different species. We show that host protease processing during viral entry is a significant barrier for several lineage B viruses and that bypassing this barrier allows several lineage B viruses to enter human cells through an unknown receptor. We also demonstrate how different lineage B viruses can recombine to gain entry into human cells and confirm that human ACE2 is the receptor for the recently emerging 2019-nCoV.

## Introduction

Severe Acute Respiratory Syndrome Coronavirus (SARS-CoV) first emerged in humans in 2003 after transmitting from animals in open air markets in China^1, 2^. Shortly thereafter, several genetically related viruses were identified in Chinese Horseshoes bats (*Rhinolophus sinicus*)^3–7^. At the same time, improvements in next generation sequencing technology lead to a boom of virus discovery, uncovering thousands of novel virus sequences in wild animal populations around the world. While most of these viruses have never been found in humans, many are genetically similar to known human viruses within the betacoronaviruses (β-CoV) genus. The β-CoVs are further divided into four lineages: lineage B, which includes SARS-CoV and the newly emerging 2019-nCoV, has approximately 200 published virus sequences whereas lineage C, which includes MERS-CoV, has over 500 viral sequences.

Every year, additional novel CoV sequences are discovered. However, there is a massive knowledge gap in the field as very little work is performed after the viral sequences are published. Therefore, it is unknown whether these novel viruses have the potential to emerge in human populations.

Current methods for studying novel β-CoVs are technically demanding. Viral isolation from field samples is rarely successful and reverse genetics recovery of recombinant virus is labor-intensive, and expensive as synthesis of a single genome can cost upwards of $15,000. These limitations are prohibitive to studying novel CoVs at the scale in which they are discovered.

Cell entry is an essential component of cross-species transmission, especially for the β-CoVs. All CoVs encode a surface glycoprotein, spike, which binds to the host-cell receptor and mediates viral entry^8^. For β-CoVs, a single region of the spike protein called the receptor binding domain (RBD) mediates the interaction with the host cell receptor. After binding the receptor, a nearby host protease cleaves the spike, which releases the spike fusion peptide, facilitating virus entry^9–12^. Known host receptors for β-CoVs include angiotensin converting enzyme 2 (ACE2) for SARS-CoV and dipeptidyl peptidase 4 (DPP4) for MERS-CoV^13, 14^.

Structural studies of coronaviruses have shown that the spike RBD is capable of folding independently from the rest of the spike protein and contains all of the structural information for host receptor binding^15^. Additionally, a previous study showed that replacing the RBD of the lineage B bat virus, Rp3, allowed the virus to enter cells expressing human ACE2 (hACE2)^16^. We therefore developed a method to functionally test the RBDs from novel lineage B β-CoVs in place of the SARS-CoV spike RBD (figure 1). Synthesizing just the RBD of spike is much faster and cost-effective than conventional pseudotyping methods that rely on synthesis of the full ∼4kb spike sequence for novel CoVs: a process that can take weeks and is cost-prohibitive for large panels of spike sequences. The short turnaround time for our approach allowed us to test the receptor usage of all published, unique RBD sequences in lineage B, and also rapidly confirm the ACE2 receptor usage of the 2019-nCoV spike, which emerged in China in January 2020 as our study was ongoing.

**Figure 1:**
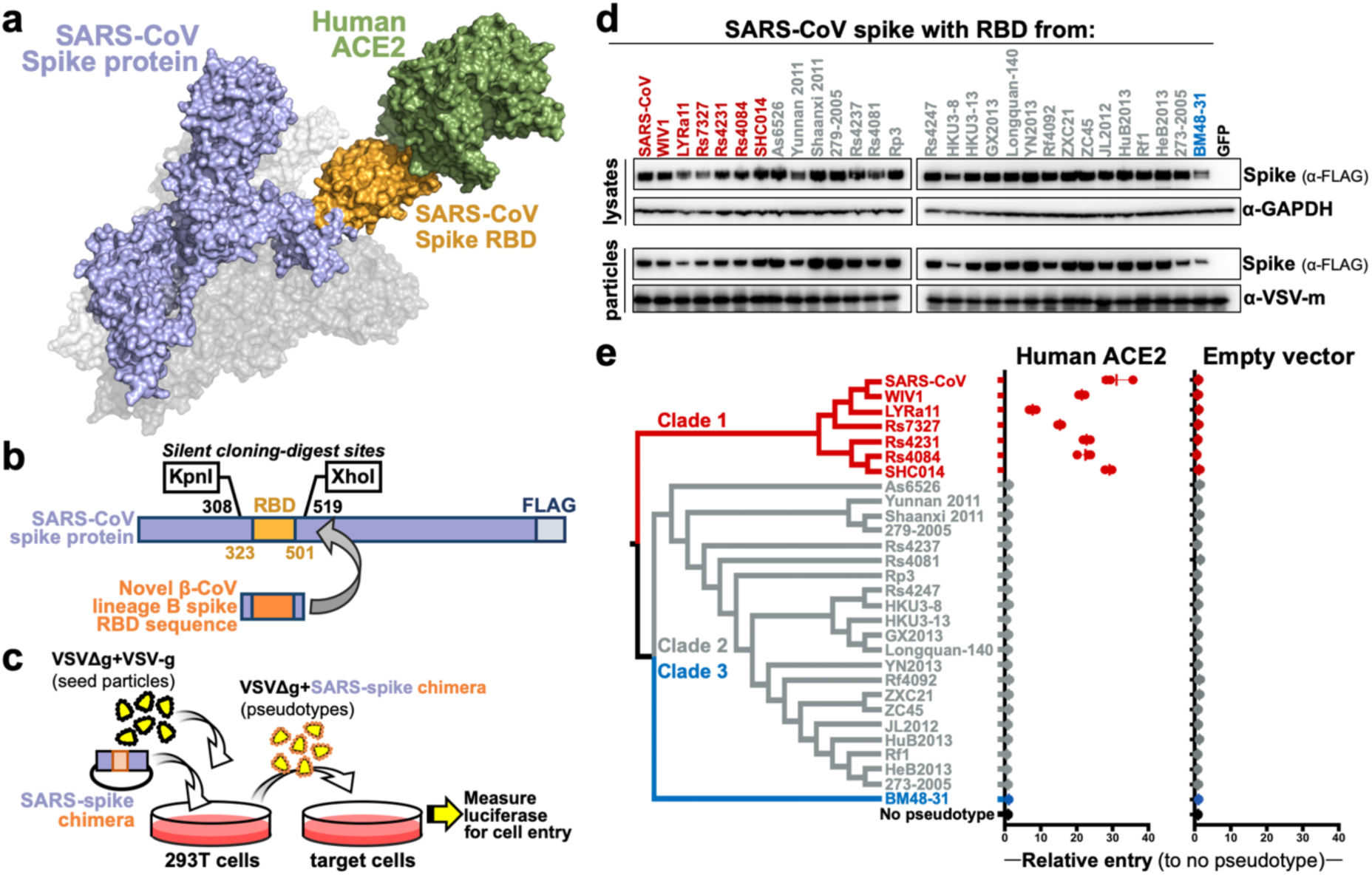
β-Coronavirus lineage B entry with human ACE2 is clade-specific. **a**, β-Coronaviruses, including SARS-CoV, interact with the host cell receptor via the Receptor Binding Domain (RBD) in spike (PDB: 5X5B, 2AJF) . **b**, SARS-CoV spike was engineered with silent mutations to facilitate cloning novel RBD sequences in place of the SARS spike RBD. SARS spike amino acid numbers are indicated for silent cloning sites and the RBD in grey and orange, respectively. **c**, outline of experimental workflow. **d**, Westernblot of producer cell lysates and concentrated reporter particles. **e**, BHK cells were transfected with either human ACE2 or empty vector, subsequently infected with VSV-reporter particles pseudotyped with chimeric spikes, luciferase was measured and normalized to no pseudotype as a readout for cell entry. Shown are the data for 3 replicates.

We show that lineage B RBDs divide into functionally distinct clades and that several previously-unappreciated viruses exhibit compatibility with an unknown receptor on human cells. We also show that these clades are capable of recombining to impart human host-cell entry phenotypes, and that, beyond the RDB-receptor interaction, host protease processing is another species barrier encountered by lineage B β-CoVs during cell entry.

## Methods

### Cells

293T, A549, BHK, Caco-2, Huh-7.5, PK-15, and Vero cells were maintained in DMEM (Sigma) supplemented with 10% FBS, penicillin/streptomycin, and L-glutamine. RhiNi/40.1, AJ-primary, AJ*i*, HypNi, RaKSM-2.5i, RhiLu, and RhiNi cells were maintained in DMEM/F12 (Gibco) supplemented with 12% FBS, penicillin/streptomycin, non-essential amino acids, sodium pyruvate and L-glutamine. AJ-primary cells were immortalized with a lentiviral vector expressing SV40 T-antigen following the manufacturer’s instructions to generate AJ*i* cells (abm; #G203). RaKSM-2.5 primary cells have been previously described and were immortalized in this study similar to AJ*i* cells^17^.

### Plasmids

The spike coding sequences for SARS-CoV Urbani, As6526, and BM48-31 were codon optimized for human cells, appended with a 5’ kozak expression sequence (GCCACC) and 3’ tetra-glycine linker followed by nucleotides encoding a FLAG tag sequence (DYKDDDDK). For SARS-CoV spike, silent mutations were introduced around codons 308 and 519 to form KpnI and XhoI digest sites. For As6526 spike, silent mutations were introduced around codons 290 and 501 to form AflII and HindIII digest sites. For BM48-31 spike, silent mutations were introduced around codons 295 and 501 to form AflII and HindIII digest sites. These engineered spike sequences were synthesized and cloned into pcDNA3.1+ (GenScript).

Spike RBDs were first codon-optimized for human cells, appended with regions of the target spike backbone to facilitate Infusion cloning and synthesized as double stranded DNA fragments (IDT DNA). SARS-CoV, As6526 or BM48-31 engineered spike plasmids were digested with their corresponding restriction enzymes and gel purified. RBD inserts were resuspended in water and Infusion cloned into gel purified, digested spike backbone vectors (Takara).

Human ACE2 (Q9BYF1.2), DPP4 (XM_005246371.3), or APN (NP_001141.2) were synthesized and cloned into pcDNA3.1+ (GenScript). All DNA constructs were verified by Sanger sequencing (ACGT Inc.).

### Receptor transfection

BHK cells were seeded in black 96-well plates and transfected the next day with 100ng plasmid DNA encoding human ACE2, DPP4, APN or empty vector, using polyethyleneimine (Polysciences). All downstream experiments were performed 24 hours post-transfection.

### Pseudotype production

Pseudotypes were produced as previously described^18^. 293T cells were seeded onto 6-well plates pre-coated with poly-L-lysine (Sigma) and transfected the next day with 1200ng empty plasmid and 400ng of plasmid encoding coronavirus spike or GFP as a no pseudotype control. Twenty-four hours later, transfected cells were infected with VSVΔG particles pseudotyped with VSV-g as previously described^19^. After one hour of incubating at 37°C, cells were washed three times and incubated in 2mL DMEM supplemented with 2% FBS, penicillin/streptomycin, and L-glutamine for 48 hours. Supernatants were collected, centrifuged at 500xG for 5 minutes, aliquoted and stored at -80.

### Luciferase-based cell entry assay

Target cells were seeded in black 96-well plates and inoculated, in triplicate, with equivalent volumes of pseudotype stocks. For trypsin experiments, pseudotype stocks were diluted 1:1 in DMEM without FBS, trypsin was added to a final concentration of 2500μg/mL and samples were incubated at 37°C for 15 minutes. Samples were then diluted again 1:1 in cold DMEM supplemented with 2% FBS and added to cells. Inoculated plates were centrifuged at 1200xG, 4°C, for 1 hour and incubated over night at 37°C. Approximately 18-20 hours post-infection, Bright-Glo luciferase reagent (Promega) was added to each well, 1:1, without removing culture media and luciferase was measured. Relative entry was calculated by normalizing the relative light unit (RLU) for spike pseudotypes to the plate RLU average for the “no pseudotype control.”

### Western blot

Producer cells (spike-transfected 293T) were lysed in 1%SDS, 150mM NaCl, 50mM Tris-HCl, 5mM EDTA and clarified by centrifugation at 14000xG for 20 minutes. Pseudotyped particles were concentrated from producer cell lysates that were overlaid a 10% OptiPrep cushion in PBS (Sigma) and centrifuged at 20,000× g for 2 hours at 4 °C. Lysates and concentrated particles were analyzed for FLAG, GAPDH and/or VSV-m expression on 10% Bis-Tris PAGE gel (ThermoFisher).

### Accession numbers

Accession numbers for all spike sequences used here can be found in figure s1b.

## Results

### ACE2 entry is lineage B clade 1-specific

The receptor binding domain (RBD) of lineage B β-CoVs is a single, continuous domain that contains all structural information necessary to interact with the host receptor (figure 1a, b). We introduced silent mutations in the codon optimized coding sequence for SARS-CoV to facilitate replacing the SARS RBD with the RBD from other lineage B viruses (figure 1b). All lineage B sequences were downloaded from online repositories and parsed to 29 unique RBD sequences, representing all published variations of the lineage B RBD (supp. fig 1a, b). The panel of 29 RBDs phylogenetically cluster into 3 clades, as previously described^5^, but these RBD clades were not apparent in phylogenetic analysis of other viral sequences, such as the RNA-dependent RNA polymerase (supp. fig 1c). All 29 RBDs were codon optimized, synthesized and cloned in place of the SARS RBD, effectively generating chimeric spike expression constructs. We then generated VSV-luciferase reporter particles pseudotyped with the chimeric spikes (figure 1c). We chose VSV over lentiviruses as our pseudotype platform because a lentiviral pseudotypes have failed to accurately reflect viral entry with novel bat coronavirus spike protein^7^. All constructs exhibited similar levels of expression in producer cells and incorporation into VSV pseudotypes, except the chimera with BM48-31 which displayed somewhat reduced expression compared to WT SARS spike (figure 1d). We then infected BHK cells expressing the receptor for SARS-CoV or empty vector (figure 1e) and observed only clade 1, which includes SARS-CoV, WIV1 and SHC014, could enter cells transfected with human ACE2 (figure 1e).

### Protease enhances clade 2 entry

After binding the host receptor, host-cell protease cleaves spike, releasing the fusion peptide and allowing for host cell entry ^20^. Previous studies have shown that absence of the host protease or incompatibility between the host protease and viral spike can block viral entry ^21–24^. To circumvent host-cell protease incompatibility or absence, we treated our lineage B pseudotype panel and infected a wide variety of cell types from different host species (figure 2, supp. fig. 2). In the absence of exogenous protease, only clade 1 infected cells from African green monkey kidney, human gastrointestinal tract, human liver, and porcine kidney, in agreement with previous studies (figure 2; supp. fig. 2a, b). Surprisingly, exogenous protease enhanced entry of a subset of clade 2 spike chimeras in nonhuman primate, bat and human cells (figure 2). Importantly, VSV-g pseudotyped particles were able to produce luciferase signal in all cell lines tested in this study (supp. fig. 2c).

**Figure 2:**
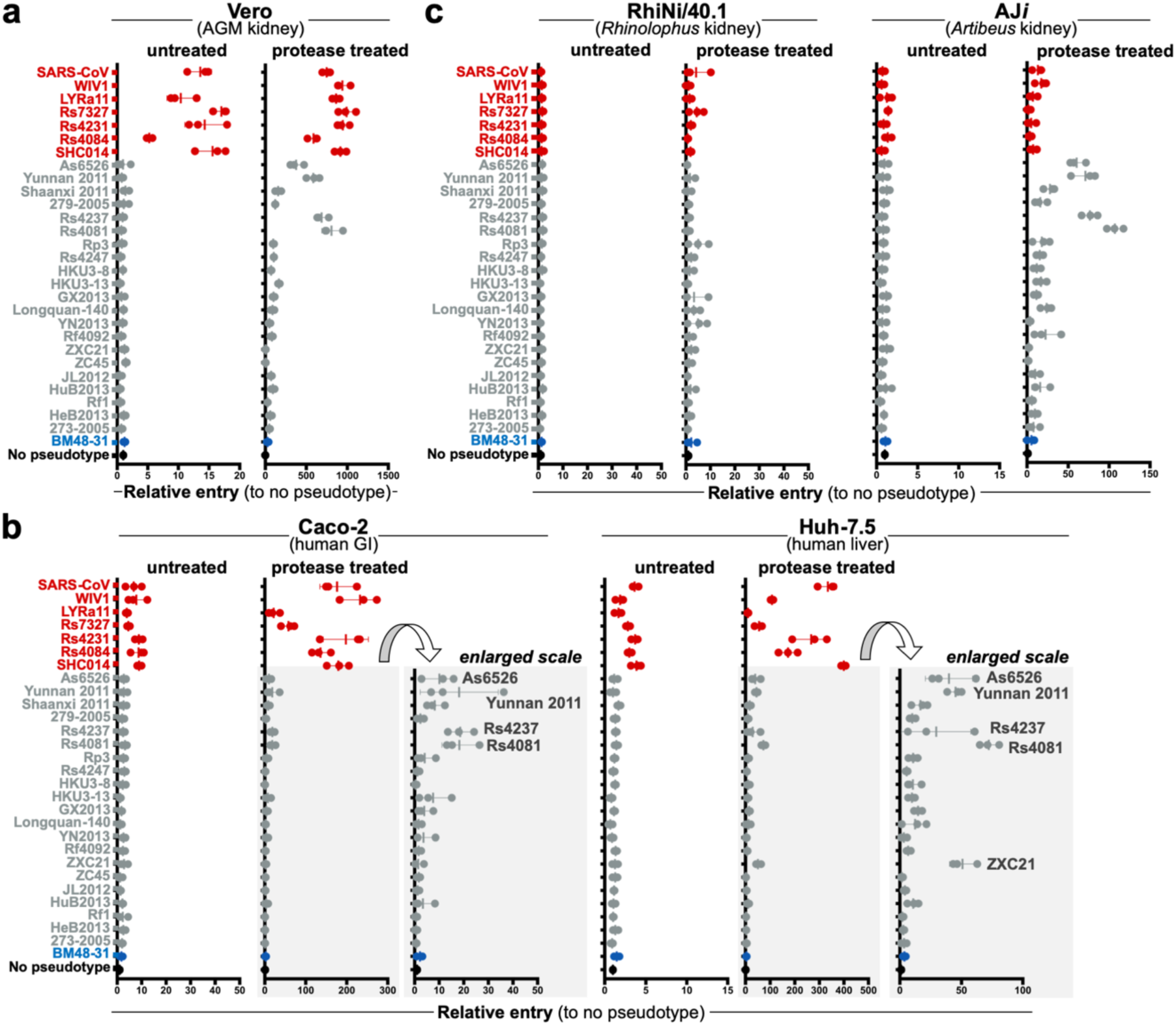
Trypsin enhances lineage B entry in various cell lines. **a**, Primate cells, **b**, human cells or **c**, bat cells were infected with VSV-particles pseudotyped with the lineage B chimeric spike panel. Pseudotypes were either left untreated or incubated with trypsin before addition to the cells. Luciferase was measured and normalized to particles produced without pseudotype. Shown are the data for 3 replicates.

### Clade 2 entry is receptor-dependent

We next tested human variants of known β-CoV receptors for their ability to mediate cell entry of clade 2 and 3 spike chimeras. We also tested human aminopeptidase N (APN), a receptor for alphacoronaviruses, which have been shown to utilize either human ACE2 or human APN for cell entry (figure 3a). Protease treatment only enhanced entry of clade 1 RBDs on cells expressing human ACE2, but not human DPP4 or APN. No entry was observed with clade 2 or 3 spikes, regardless of receptor or protease addition. Human dipeptidyl peptidase IV (DPP4), the receptor for the lineage C β-CoVs, MERS-CoV, only mediated entry of MERS-CoV (figure 2b, middle panels). Importantly, in the absence of receptor, no entry was observed for any of the pseudotypes, suggesting that protease-mediated entry is receptor-dependent (figure 2b, right panels).

**Figure 3:**
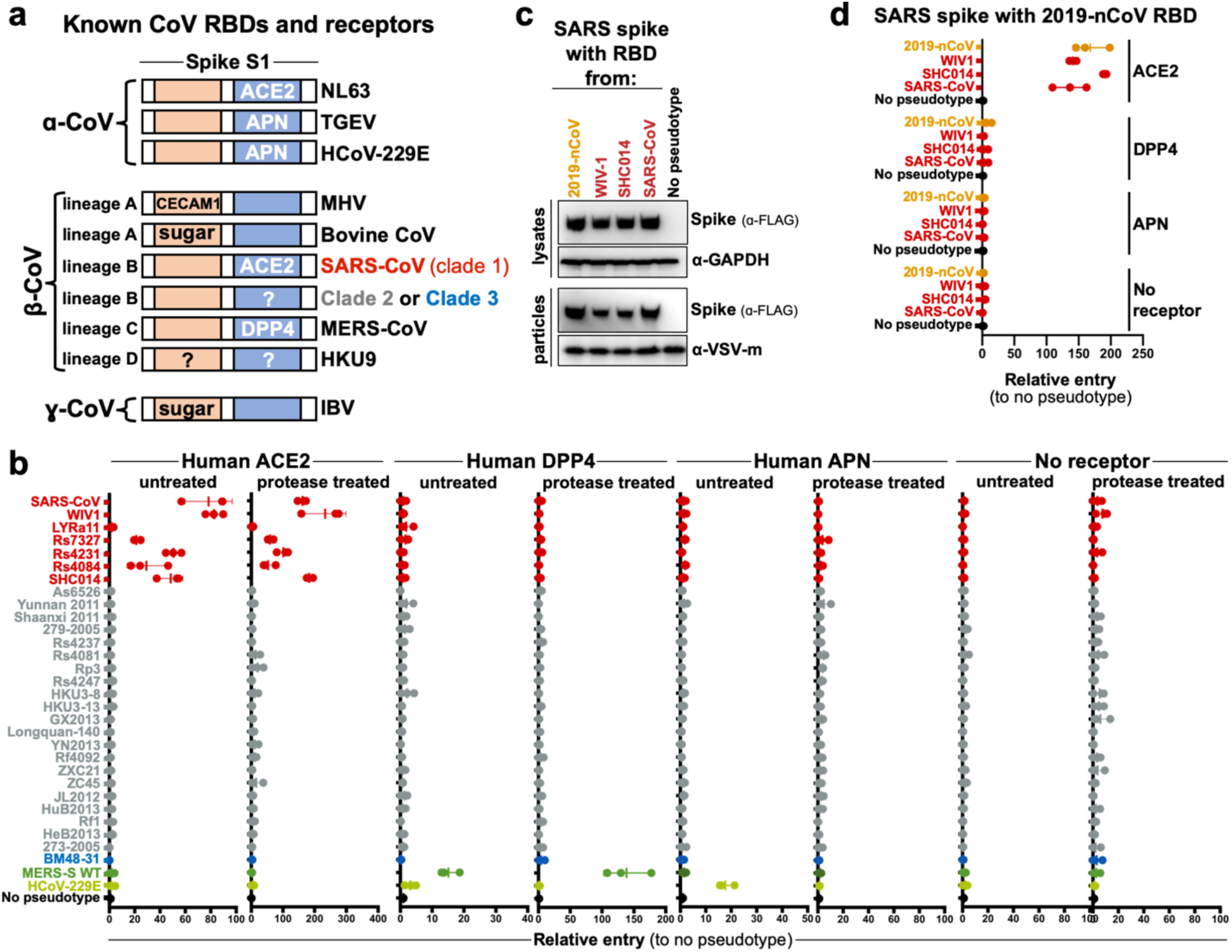
Lineage B entry into cells with known CoV receptors. **a**, Schematic of known coronavirus RBDs and their receptors. **b**, Pseudotyped particles were either left untreated and treated with trypsin and subsequently used to infect BHK cells transfected with the coronavirus receptor indicated. Shown are data from 3 replicates. **c**, Expression and pseudotype incorporation of SARS-S-2019-nCoV RBD chimeras. **d**, Pseudotypes were used to infect cells expressing hACE2, hDPP4, hAPN, or empty vector, without protease treatment.

### Receptor usage of 2019-nCoV

While our study was ongoing, a novel lineage B virus tentatively named 2019- nCoV was identified as the cause of a pneumonia outbreak in Hubei, China. Once the sequence was publicly available, we synthesized, cloned and tested the RBD from 2019- nCoV in our assay with human variants of known coronavirus receptors. The chimeric SARS-2019-nCoV spike protein expressed and was incorporated into particles similarly to other clade 1 chimeric spikes (figure 3c). The 2019-nCoV RBD was capable of entering cells expressing human ACE2, but not any of the other receptors tested (figure 3d; s3).

### Clade determinants for ACE2 usage

Consensus sequences of the three lineage B clades showed several key differences between these groups. Only clade 1 RBDs contain all 14 residues that have been shown through crystallography, to interact with human ACE2 (figure 4a; s4). The majority of these residues are absent from clades 2 and 3, which contain additional deletions in surface exposed loops that cluster at the interface with ACE2 (figure 4 a, b). We generated a series of clade consensus RBD variants to determine the minimum number of mutations needed to impart ACE2 function on clade 2 and 3 RBDs (figure 4c). Introducing the two loop deletions from clade 1 in clade 2 results in a reduced spike expression, impaired pseudotype incorporation and loss of cell entry (figure 4c, d). Restoring these loops in clade 2 and 3 from the loops found in clade 1 did not enhance entry with ACE2 (figure 4c; 2→1 and 3→1 version 1). Introducing all 14 ACE2 contact points in clade 2 or 3 also failed to restore ACE2 entry (figure 4c; 2→1 and 3→1 version 2). Only replacing all 14 contact points and the surrounding amino acids (also known as the receptor binding motif, RBM) lead to increased ACE2 entry with clade 2 and 3 RBDs (figure 4c; 2→1 version 3 = clade 2 residues 322-400 + clade 1 residues 400-501; 3→1 version 3 = clade 3 residues 322-385 + clade 1 residues 386-501). Taken together, these results show that the entire RBM from clade 1 is needed for ACE2 entry.

**Figure 4:**
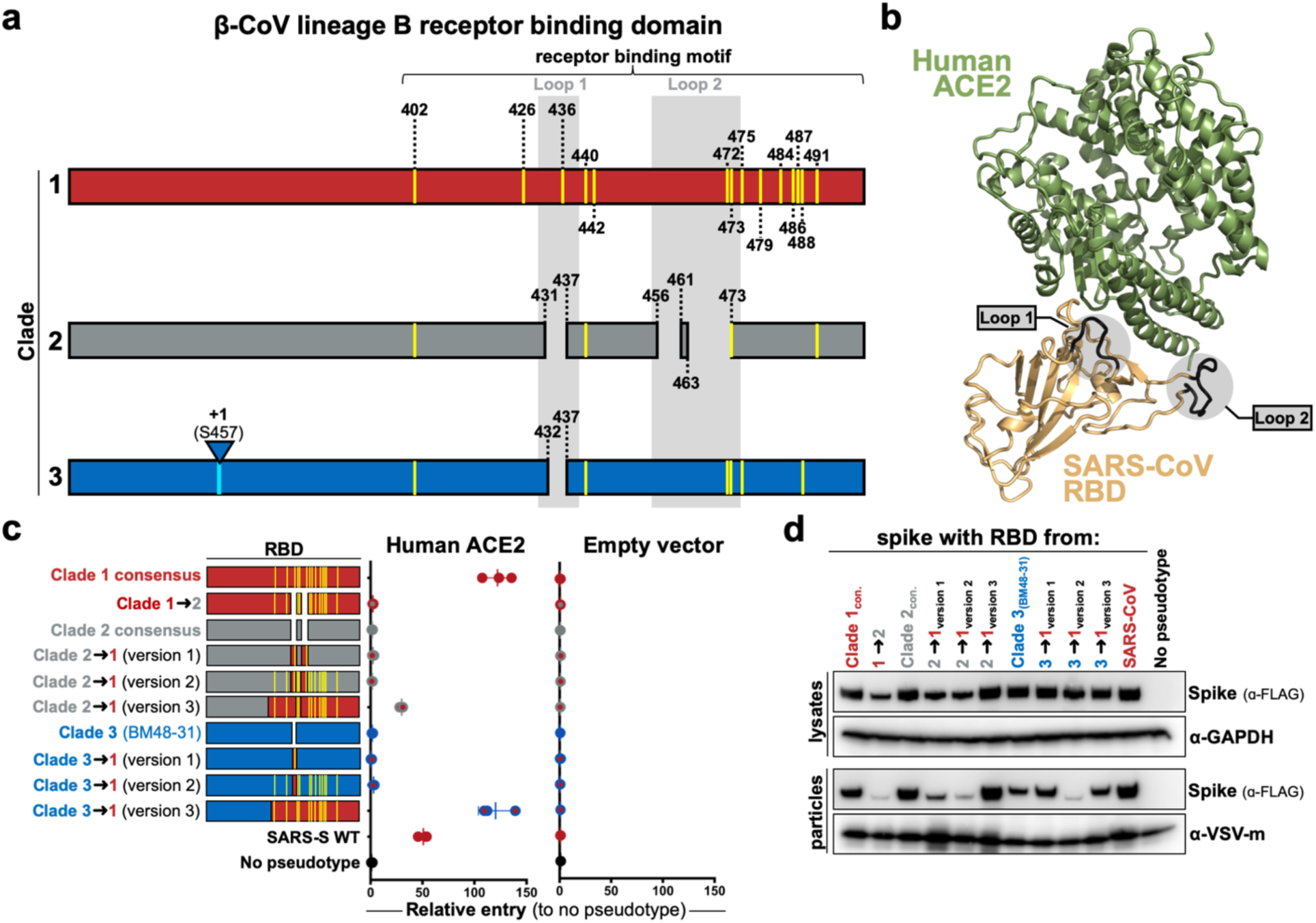
Lineage B clade-specific determinants for human ACE2 usage. **a**, Schematic overview of clade 1, 2 and 3. Highlighted in yellow are the 14 residues that contact human ACE2. Deletions in loops 1 and 2 are indicated for clades 2 and 3. **b**, Structure of human ACE2 and the SARS-S RBD (PDB: 2AJF), with loops highlighted in gray. **c**, VSV pseudotypes were generated with the indicated RBD and used to infect BHKs transfected with either human ACE2 or empty vector. Shown are data for 3 replicates. **d**, Westernblot of producer cell lysates and concentrated pseudotyped particles.

### Full-spike and RBD chimeras are comparable

We next synthesized full-length clade 2 and 3 spikes to compare to our RBD chimeras. We selected the clade 2 spike, As6526, because it consistently gave strong entry signal in human cells following protease-treatment (figure 2b) as well as BM48-31, the only clade 3 spike in our panel. As we did for SARS-CoV spike, clade 2 and 3 spikes were codon optimized, FLAG-tagged and silent mutations were introduced to facilitate replacing their RBD with the consensus RBD from clade 1 (figure 5a). All chimeric constructs expressed similarly, with the exception for the SARS-BM48-31 RBD chimera, which exhibited reduced expression and incorporation (figure 5b). Protease treatment enhanced entry of both the As6526 clade 2 RBD chimera and full-length spike entry into Huh cells (figure 5c). Protease treatment had no effect on either the BM48-31 clade 3 chimera or full-length spike (figure 5c). Taken together, these findings show that SARS- lineage B RBD chimeras reflect the entry phenotype of full-length lineage B spikes. Finally, we tested if receptor-binding and protease processing are coupled. We replaced the RBD of full-length clade 2 and 3 spike with the consensus RBD from clade

**Figure 5:**
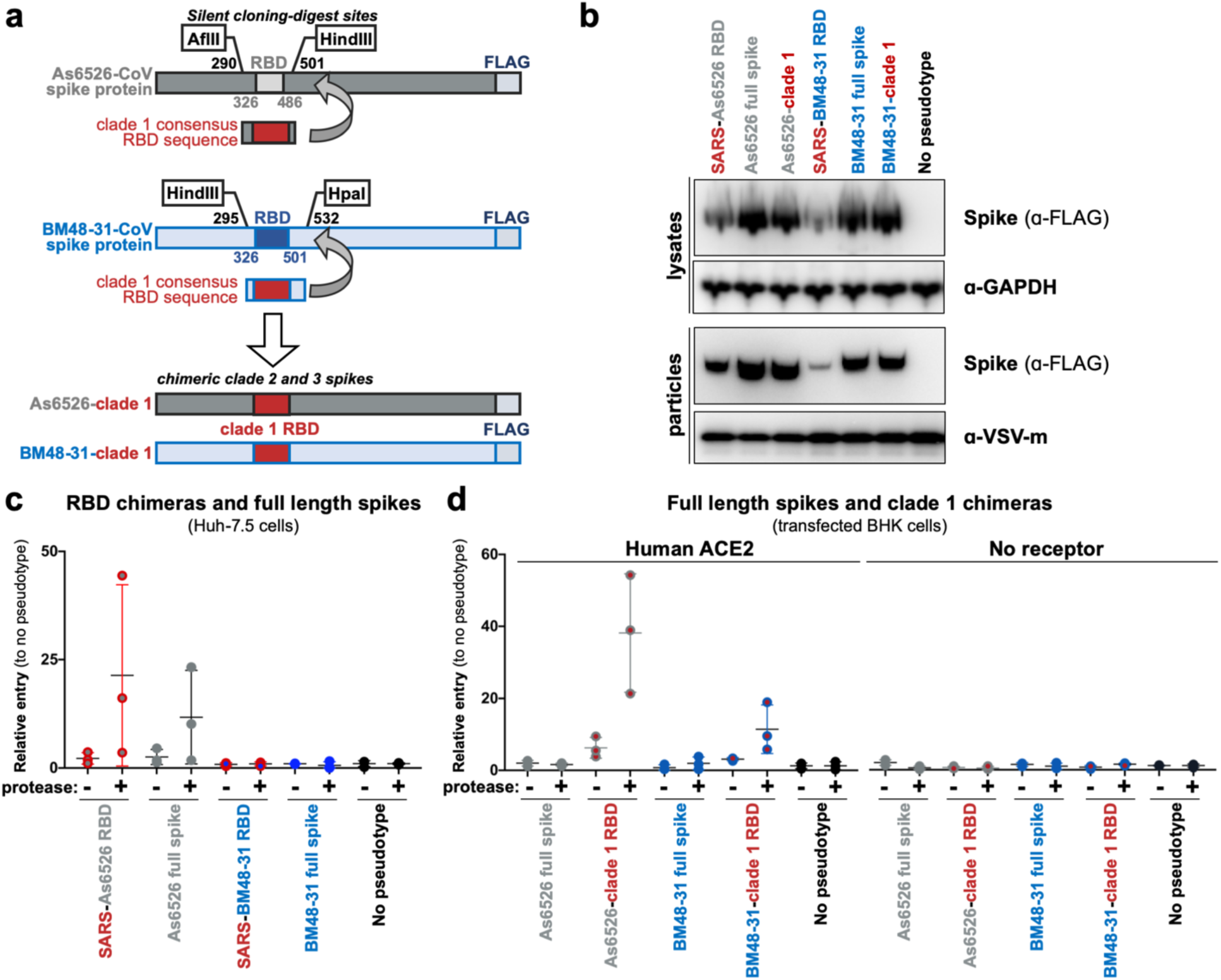
Comparison of chimeric and full-length clade 2 and 3 spikes. **a**, Full length spike sequences from As6526 (clade 2) and BM48-31 (clade 3) were codon optimized, FLAG tagged and synthesized. Silent mutations flanking the RBD facilitated replacing the native RBD with the clade 1 consensus RBD. **b**, Westernblot of producer cell lysates and concentrated pseudotypes particles. **c**, Pseudotypes with indicated spike constructs were left untreated or treated with trypsin and subsequently used to infect Huh-7.5 cells. Shown are data for 3 replicates. **d**, Pseudotypes with indicated spike constructs were left untreated or treated with trypsin and subsequently used to infect BHK cells transfected with human ACE2. Shown are data for 3 replicates.

1 and tested pseudotypes on cells expressing the clade 1 receptor. The clade 1 consensus RBD efficiently facilitated entry of both As6526 and BM48-31 spike only following protease treatment. These findings show that even though BHK-hACE2 cells support full-length clade 1 spike entry, just having the RBD from a clade 1 virus is insufficient to mediate entry. As seen in our previous experiments, protease treatment did not enhance pseudotype entry in the absence of host receptor (figure 5).

## Discussion

Despite significant advances in next generation sequencing technologies, which have facilitated the discovery of thousands of novel animal-derived viruses, tools for downstream functional assessment of these novel sequences are lacking. To gain traction on this ever-growing problem, we took a reductionist approach to coronavirus entry and developed a scalable, BSL2-compatible method for testing only the minimal region of the virus essential for interacting with the host receptor (figure 1, figure s1a). Because most of these viruses have never been isolated, we resorted to synthetic biology and molecular engineering to reduce the burden of gene synthesis to just a small fragment. Thus, the cost and synthesis production time for testing several spikes for entry in our system is dramatically reduced (figure s1d). In theory, this approach to functional viromics should be applicable to a wide variety of virus-host proteins and interactions.

Coronavirus entry is a multi-step process involving multiple, distinct domains in spike that mediate virus attachment to the cell surface, receptor engagement, protease processing and membrane fusion^8^. While the RBD:receptor interaction is most studied in this process, recent studies have highlighted the major role host protease processing plays as a species barrier^22, 25–27^. Lineage C coronaviruses include MERS-CoV as well as distantly related viruses such as HKU4, HKU5 and PDF-2180 ^28, 29^. Studies have shown that HKU4 can bind human DPP4 but requires addition of exogenous trypsin to facilitate cell entry and that HKU5 and PDF-2180 spikes can enter human cells through an unidentified receptor with protease treatment^22, 26^. Analogous to these earlier studies of lineage C CoVs, we observed protease-enhanced entry of lineage B CoVs (figure 2, 3, 5). While it has been shown that host proteases cleave spike, allowing for downstream membrane fusion, other evidence suggests that protease may act on the receptor as well to activate it^30^. Addition of protease during the course of SARS-CoV infection facilitated entry in cells with low-expression of ACE2 that is normally insufficient to support virus entry^30^. Indeed, we saw evidence of residual trypsin activity on the cells after infection in our studies as the cell monolayer was loose compared to the untreated condition. Similarly, Menachery et al. also observed cell rounding during their trypsin infections^26^. Therefore, further studies are needed to assess where trypsin is enhancing entry of coronaviruses: at the level of spike, the receptor, or both.

In the absence of exogenous protease, only clade 1 RBDs entered nonhuman primate, human and porcine cell lines (figure 2a, b, s2a, b). These findings are in strong agreement with previous studies that have either isolated virus (WIV1) or rescued recombinant chimeric viruses (SHC014, Rs4231, Rs7237)^5, 7, 31^. However, with trypsin, a subset of genetically-similar clade 2 RBDs gained entry in these cells, suggesting their barrier is at the level of protease processing (figure 2a, b). The other spike from clade 2 and 3 did not enter the cells we tested, regardless of protease addition, suggesting an absent or incompatible receptor. Surprisingly, the protease-dependent entry phenotype was consistent in the reverse spike chimeras in which we replaced the RBD in clade 2 or 3 spike with a clade 1 RBD (figure 5d), suggesting that either the protease site between S1/S2 is not compatible with the chimeric spike backbone or the protease is not expressed in these cells (figure s5). Because clade 1 spikes enter cells expressing human ACE2 without addition of protease but clade2-clade1 chimeras require protease, our data suggests the spike protease cleavage site is adapted to the protease environment of the receptor-bound RBD (figure s5).

None of the spike pseudotypes efficiently entered *Rhinolophus* cells, which has been observed in previous studies using these cells^32, 33^ (figure 2c). Surprisingly, AJi cells were selectively permissive for only clade 2 entry following protease treatment, which suggests that clade 2 RBDs interact with a receptor that is distinct from clade 1 (figure 2c).

Our results show that, despite all being classified as the same virus species, most lineage B β-CoVs do not use currently known coronavirus receptors (figure 1e, 3a, b). Critically, we did not observe any pseudotype entry in the presence of protease and absence of receptor, suggesting that lineage B cell entry is still receptor-dependent following protease treatment (figure 3b). While our study was ongoing, a novel lineage B β-CoV was identified as the etiological agent behind an outbreak of pneumonia in Wuhan, Hubei, China (2019-nCoV). The RBD for 2019-nCoV has residues and motifs found in all 3 clades but forms a distinct clade, so we tested it for receptor usage and observed entry only with human ACE2 but not other known coronavirus receptors (figure 3d). Interestingly, within the backbone of SARS-CoV spike, cell entry of 2019-nCoV was similar to the other clade 1 spikes tested, including SARS-CoV. These finding suggests 2019-nCoV is capable of using human ACE2 as efficiently as SARS-CoV, which may help explain the human-to-human transmissibility of this virus. More studies are needed with the full spike sequence and, ideally, a viral isolate.

The receptor binding motif (RBM) is a small region in the C-terminal half of the RBD and contains all the residues that interface with the host receptor (figure 3a)^15^. The 14 contact points in the co-structure of the SARS-RBD bound to human ACE2 are largely absent from clade 2 and 3 RBDs, which also contain deletions compared to clade 1 RBMs (figure 4a, b, s4a). Simply mutating clade 2 and 3 to have the 14 contact points was insufficient to impart human ACE2 usage (figure 4c). This is likely because the non-contact residues in the RBM play a supportive and structural role for these contact points, and indeed, these non-contact residues are different between the clades (figure s4).

In contrast to changing individual amino acids, our chimeric RBD constructs show that clade 2 and 3 RBD containing the clade 1 RBM are compatible with human ACE2. Coronaviruses frequently undergo recombination, gaining large swaths of genetic material at once ^34, 35^. Taken together with our data, it is possible that recombination with a clade 1 virus will impart compatibility with human ACE2. Interestingly, the 2019-nCoV RBD forms a clade that is distinct from the other 3 clades (figure s1c). However, the 2019- nCoV RBD contains most of the contact points with human ACE2 that are found in clade 1 as well as some amino acid variations that are unique to clade 2 and 3 (figure s4b). Taken together with our receptor assay results, it may be possible that 2019-nCoV arose from recombination between clade 1 and the other clades.

As we saw with the SARS-As6526 RBD (clade 2) spike chimera, full length As6526 spike entered cells following protease treatment, but BM48-31 (clade 3) spike did not (figure 2, 5c). These data show that the chimeric spikes generally reproduce the entry phenotypes of full-length spikes. Notably, the full length As6526 spike did not enter cells as efficiently as the SARS-As6526 chimera, suggesting that other human-cell adaptations are likely needed in As6526 spike.

The capacity to predict the zoonotic potential of newly detected viruses has been severely hindered by a lack of functional data for these novel animal virus sequences. Here, we have developed a rapid and cost-effective platform to functionally test large groups of related viruses for zoonotic potential. We found that several other lineage B coronaviruses are capable of entering human cells through an unknown receptor and that lineage B spike proteins can recombine to gain entry with a known host-receptor. Taken together with the latest outbreak of 2019-nCoV in humans, these findings underscore the importance of continued surveillance of coronaviruses at the sequence and functional levels in order to better prepare for the next emerging virus.

## Acknowledgements

We would like to thank Dr. Anthony Schountz for generously sharing primary artibeus cells, as well as Dr. Marcel Muller for providing the *Rhinolophus* cells used in this study. We would also like to acknowledge Wu,F., Zhao,S., Yu,B., Chen,Y.-M., Wang,W., Hu,Y., Song,Z.-G., Tao,Z.-W., Tian,J.-H., Pei,Y.-Y., Yuan,M.L., Zhang,Y.-L., Dai,F.-H., Liu,Y., Wang,Q.-M., Zheng,J.-J., Xu,L., Holmes,E.C. and Zhang,Y.-Z, the Wuhan Institute of Virology, Chinese Academy of Medical Sciences & Peking Union Medical College, Chinese Center for Disease Control and Prevention, and Wuhan Jinyintan Hospitar for providing the genome sequence for 2019-nCoV and other essential outbreak information to the scientific community. This work was supported by the Intramural Research Program of the National Institute of Allergy and Infectious Diseases.

