## Supplemental figures for "Functional assessment of cell entry and receptor usage for lineage B β-coronaviruses, including 2019-nCoV"

### SUPPLEMENTAL INFORMATION

### Corresponding authors:

Drs. Vincent Munster and Michael Letko

Rocky Mountain Laboratories

NIAID/NIH

903S 4th Street

Hamilton, MT 59840

**a**

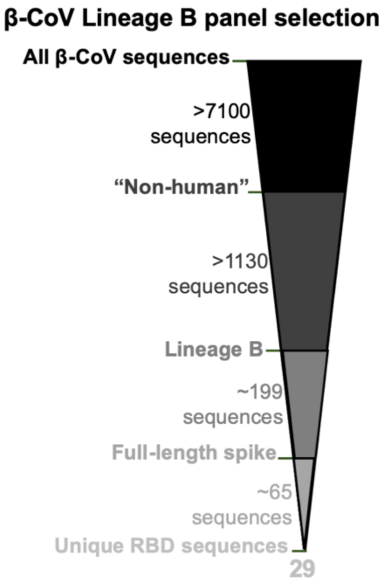


**b**

| | Virus | Accession | Clade | Host species | Location |
|---|---|---|---|---|---|
| 1 | SARS Urbani | AY278741 | 1 | human | Guangdong, China (origin of outbreak) |
| 2 | WIV1 | KF367457 | 1 | *Rhinolophus sinicus* | Yunnan, China |
| 3 | LYRa11 | KF569996 | 1 | *Rhinolophus affinis* | Baoshan, Yunnan, China |
| 4 | Rs7327 | KY417151 | 1 | *Rhinolophus sinicus* | Kunming, Yunnan Province, China |
| 5 | Rs4231 | KY417146 | 1 | *Rhinolophus sinicus* | Kunming, Yunnan Province, China |
| 6 | Rs4084 | KY417144 | 1 | *Rhinolophus sinicus* | Kunming, Yunnan Province, China |
| 7 | RsSHC014 | KC881005 | 1 | *Rhinolophus sinicus* | Yunnan, China |
| 8 | Wuhan-CoV | MN908947 | 1/2 | human | Wuhan, Hubei, China |
| 9 | As6526 | KY417142 | 2 | *Aselliscus stoliczkanus* | Kunming, Yunnan Province, China |
| 10 | Yunnan2011 | JX993988 | 2 | *Chaerephon plicata* | Yunnan, China |
| 11 | Shaanxi2011 | JX993987 | 2 | *Rhinolophus pusillus* | Shaanxi, China |
| 12 | 279-2005 | DQ648857 | 2 | *Rhinolophus macrotis* | Hubei, China |
| 13 | Rs4237 | KY417147 | 2 | *Rhinolophus sinicus* | Kunming, Yunnan Province, China |
| 14 | Rs4081 | KY417143 | 2 | *Rhinolophus sinicus* | Kunming, Yunnan Province, China |
| 15 | Rp3 | DQ071615 | 2 | *Rhinolophus pearsoni* | Nanning, Guangxi, China |
| 16 | Rs4247 | KY417148 | 2 | *Rhinolophus sinicus* | Kunming, Yunnan Province, China |
| 17 | HKU3-8 | GQ153543 | 2 | *Rhinolophus sinicus* | Guangdong, China |
| 18 | HKU3-13 | GQ153548 | 2 | *Rhinolophus sinicus* | Guangdong, China |
| 19 | GX2013 | KJ473815 | 2 | *Rhinolophus sinicus* | Guangxi, China |
| 20 | Longquan-140 | KF294457 | 2 | *Rhinolophus monoceros* | Longquan, Zhejiang, China |
| 21 | YN2013 | KJ473816 | 2 | *Rhinolophus sinicus* | Yunnan, China |
| 22 | Rf4092 | KY417145 | 2 | *Rhinolophus ferrumequium* | Kunming, Yunnan Province, China |
| 23 | ZXC21 | MG772934 | 2 | *Rhinolophus sinicus* | Zhoushan City, Zhejiang, China |
| 24 | ZC45 | MG772933 | 2 | *Rhinolophus sinicus* | Zhoushan City, Zhejiang, China |
| 25 | JL2012 | KJ473811 | 2 | *Rhinolophus ferrumequium* | Jilin, China |
| 26 | HuB2013 | KJ473814 | 2 | *Rhinolophus sinicus* | Hubei, China |
| 27 | Rf1 | DQ412042 | 2 | *Rhinolophus ferrumequium* | Yichang, Hubei, China |
| 28 | HeB2013 | KJ473812 | 2 | *Rhinolophus ferrumequium* | Hebei, China |
| 29 | 273-2005 | DQ648856 | 2 | *Rhinolophus ferrumequium* | Hubei, China |
| 30 | BM48-31 | NC014470 | 3 | *Rhinolophus blasii* | Strandja Nature Park, Bulgaira |

**c** **d**

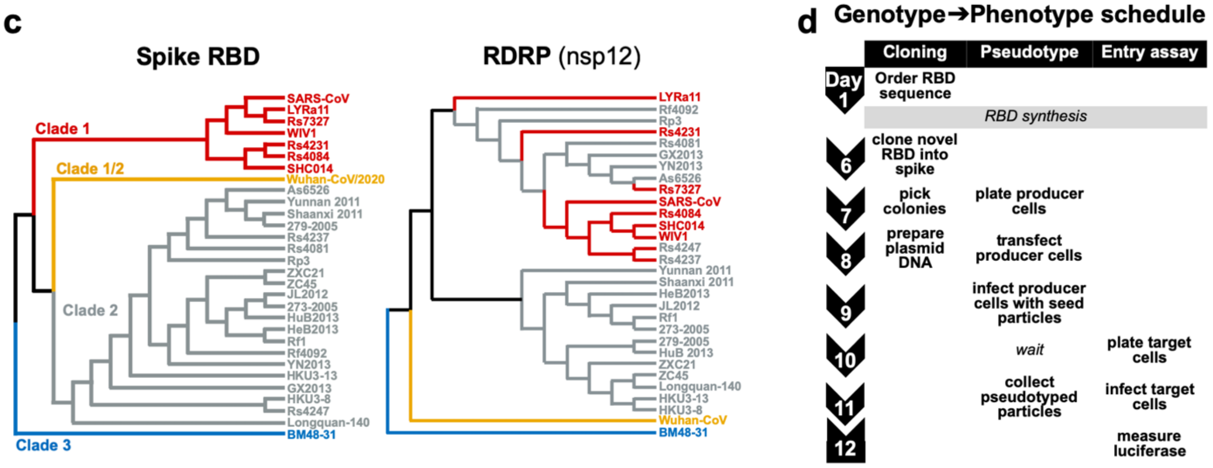


**Supplemental figure 1: Lineage B RBD panel assembly and phylogenetic analysis**
**a,** Coronavirus sequences were downloaded from NCBI and further parsed to 29 unique RBD variants. **b,** Virus isolate name, accession number, clade, host species and location of identification listed for the 29 unique lineage B RBDs used in this study. **c,** Cladograms for the spike RBD and coronavirus RNA-dependent RNA polymerase (nsp12). **d,** Overview of experimental timeline.

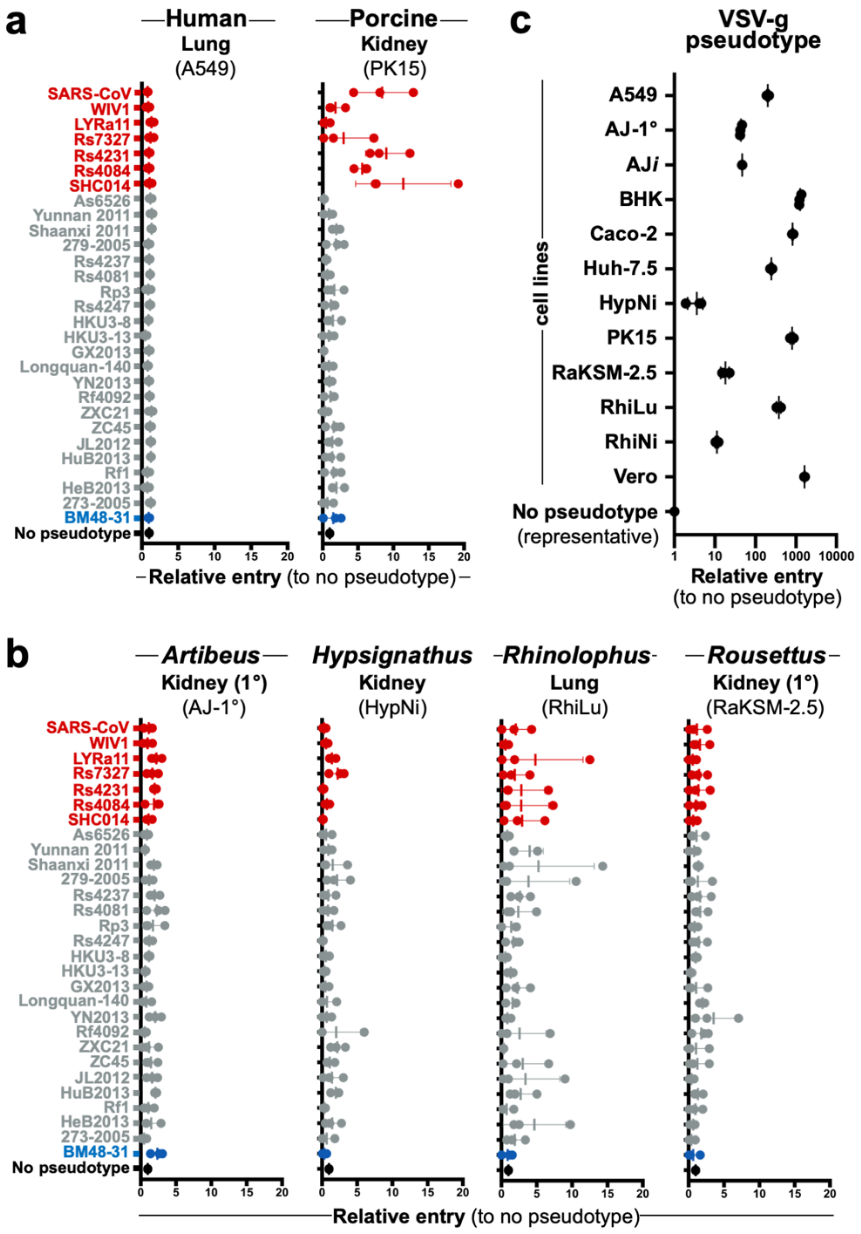


**Supplemental figure 2: additional cell lines tested without protease**

**a,** Additional human and cell lines or **b,** bat-derived cell lines derived from other species were infected with VSV-reporter particles pseudotyped with chimeric spikes and luciferase was measured a readout for cell entry. Trypsin was not used in these infections. **c,** All cell lines tested in this study supported entry and reporter expression of VSV-g pseudotyped particles. Shown are the data for 3 replicates.

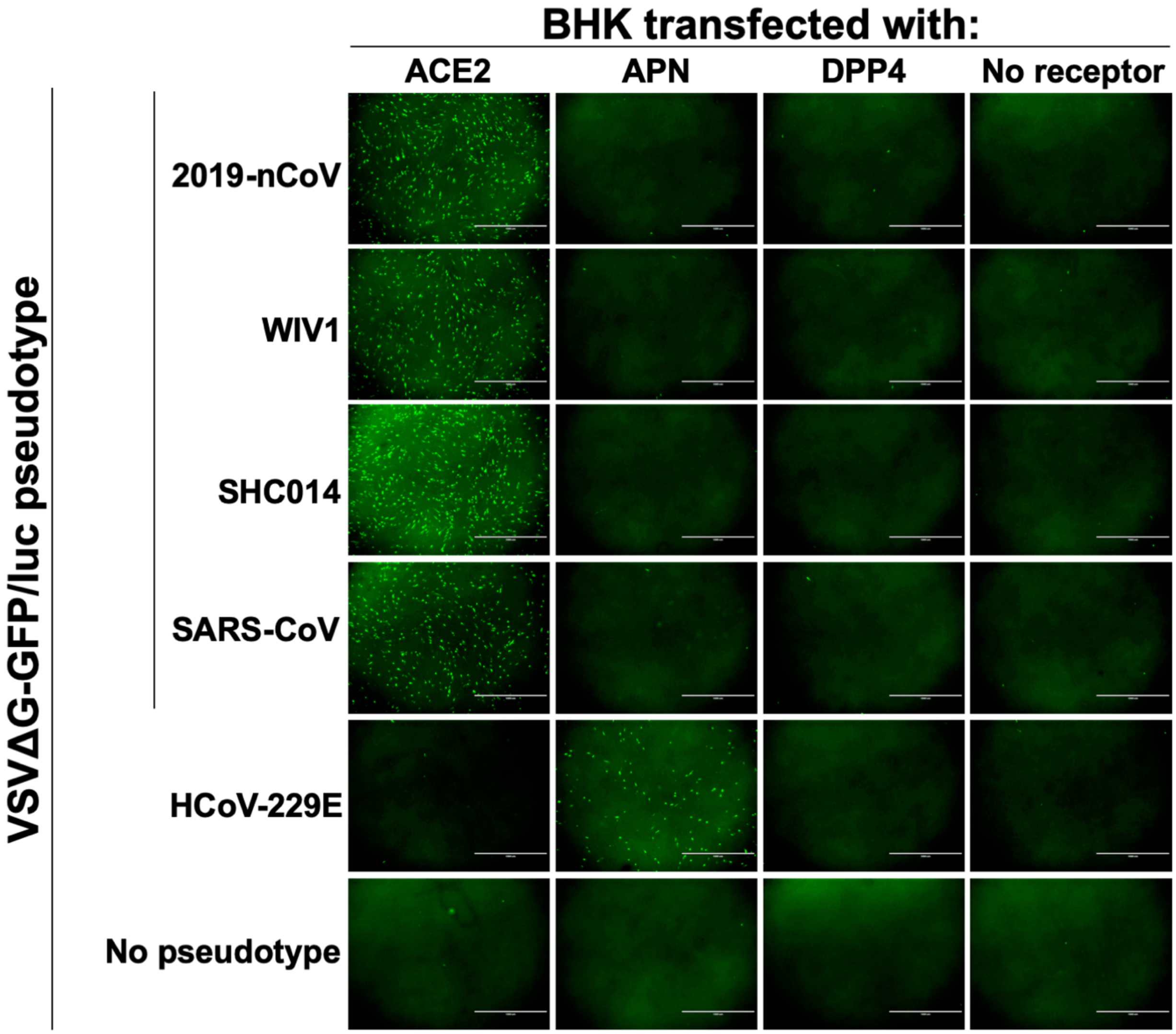


**Supplemental figure 3: 2019-nCoV uses human ACE2 to enter cells**

VSVΔG-GFP/luciferase particles were pseudotyped with the indicated spikes and used to infect BHKs transfected with known coronavirus receptors. Microscopy images were taken 12-20 hours-post infection.

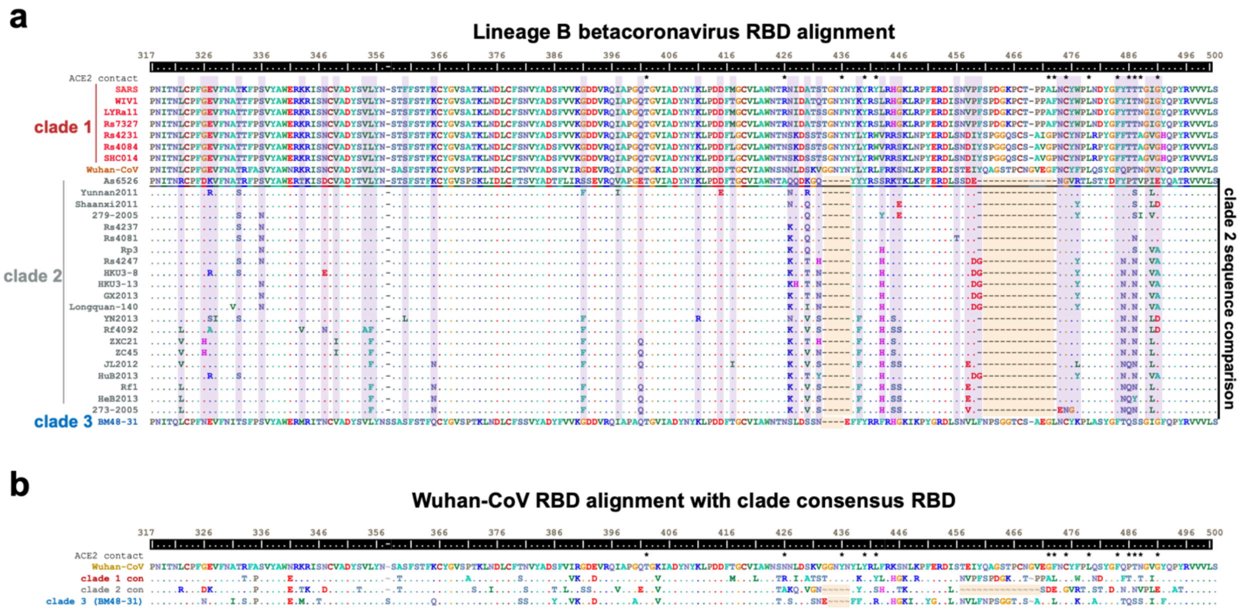


**Supplemental figure 4: Lineage B panel RBD sequence features**

**a,** Amino acid sequences corresponding to SARS-spike residues 317 through 500 were aligned with ClustalW. Contact points between SARS-spike and human ACE2 are indicated with an (*). Clade 2 sequences are shown as compared to clade 2 As6526, with identical residues indicated with a (.) and sites that vary between clade 2 viruses highlighted in purple. Loop deletions are highlighted in orange. **b,** Amino acid alignment of 2019-nCoV RBD and consensus RBD sequences for clade 1 and 2 and BM48-31 (clade 3). Loop deletions are highlighted in orange.

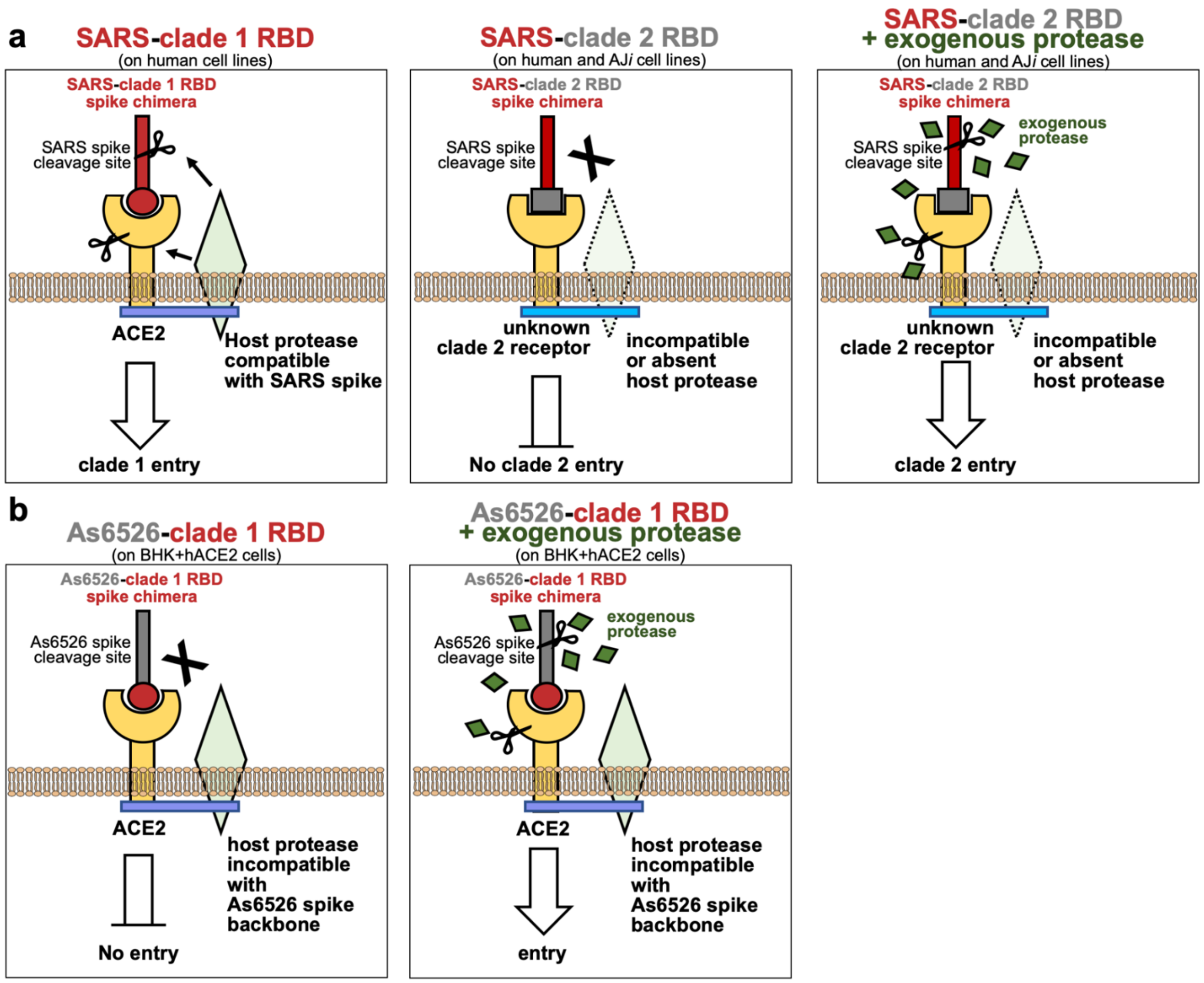


**Supplemental figure 5: model of lineage B entry**

**a,** SARS spike with a clade 1 RBD enters cells expressing ACE2 and a host protease capable of cleaving SARS spike (left panel). While the clade 2 RBD can bind an unknown host receptor, the SARS spike backbone is incompatible with the receptor-associated protease, resulting in a lack of cleavage and entry (middle panel). The addition of exogenous protease may overcome the lack of endogenous protease cleavage of spike, resulting in receptor-dependent entry. Alternatively, the addition of exogenous protease may activate the receptor to facilitate entry. **b,** Replacing the RBD of As6526 spike with the clade 1 RBD allows for ACE2 interaction, but the As6526 spike backbone is incompatible with the ACE2-associated protease (left panel). Addition of exogenous protease overcomes protease incompatibility, allowing for ACE2-mediate entry (right panel).
